# Ultra-pure isolation of low density neutrophils casts doubt on their exceptionality in health and disease

**DOI:** 10.1101/2020.06.17.156588

**Authors:** Gareth R Hardisty, Frances LLanwarne, Danielle Minns, Jonathan L Gillan, Donald J Davidson, Emily Gwyer Findlay, Robert D Gray

## Abstract

Low density neutrophils (LDNs) are described in a number of inflammatory conditions, cancers and infections and associated with immunopathology, and a mechanistic role in disease. The role of LDNs at homeostasis in healthy individuals has not been investigated. We have developed an isolation protocol that generates high purity LDNs from healthy donors. Healthy LDNs were identical to healthy NDNs, aside from reduced neutrophil extracellular trap formation. CD66b, CD16, CD15, CD10, CD54, CD62L, CXCR2, CD47 and CD11b were expressed at equivalent levels in LDNs and normal density neutrophils (NDNs) and underwent apoptosis and ROS production interchangeably. Healthy LDNs had no differential effect on CD4^+^ or CD8^+^ T cell proliferation or IFNγ production compared with NDNs. LDNs were generated from healthy NDNs *in vitro* by activation with TNF, LPS or fMLF, suggesting a mechanism of LDN generation in disease however, we show neutrophilia in people with Cystic Fibrosis (CF) was not due to increased LDNs. LDNs are present in the neutrophil pool at homeostasis and have limited functional differences to NDNs. We conclude that increased LDN numbers in disease reflect the specific pathology or inflammatory environment and that neutrophil density alone is inadequate to classify discrete functional populations of neutrophils.

**Graphical abstract:** 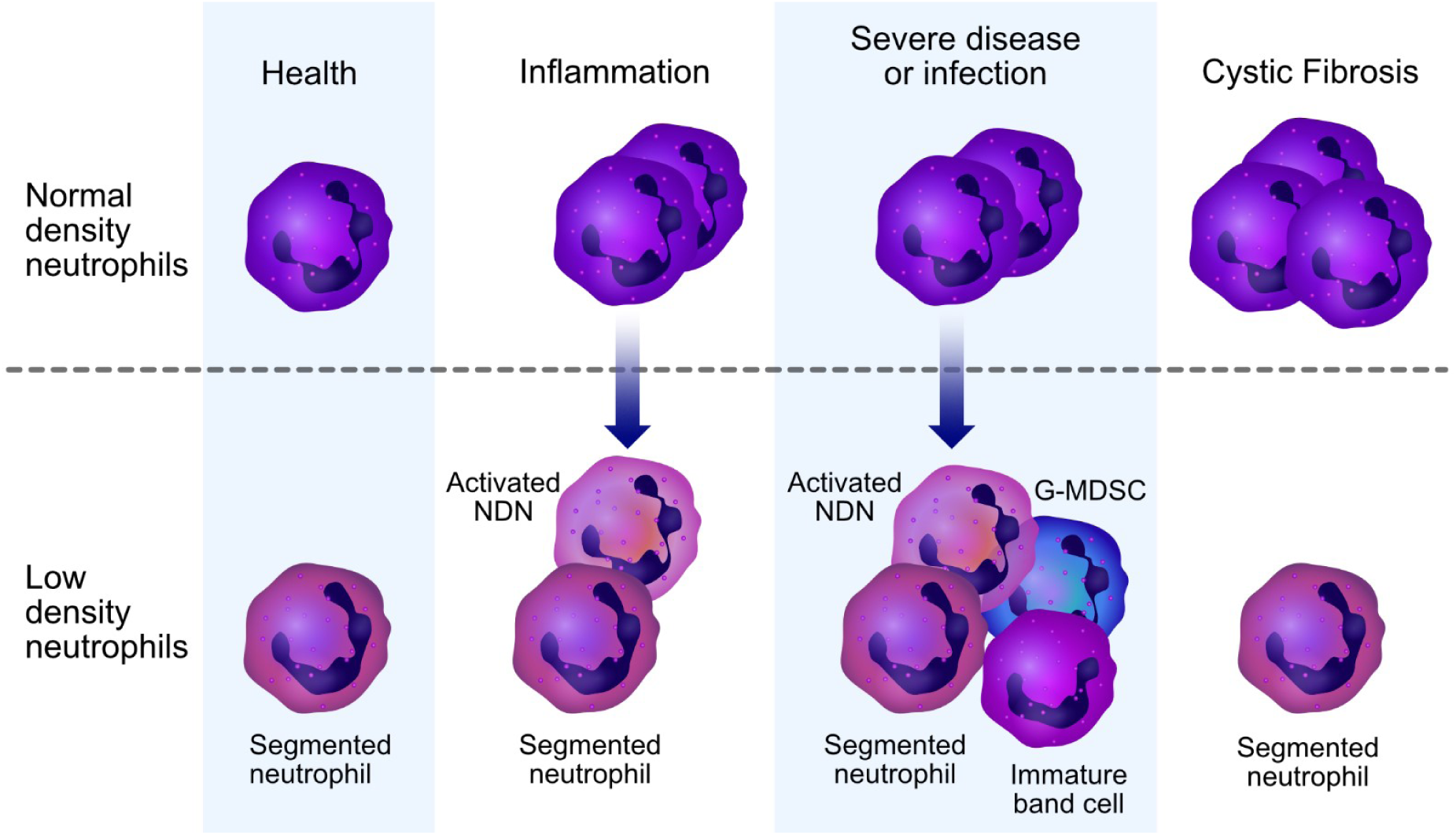

## Introduction

Heterogeneity is a characteristic of human neutrophil populations,[1] yet whether this represents truly diverse neutrophil subsets, plasticity during the immune response or differences in maturation state is undetermined. Low density neutrophils (LDNs) are a suggested subpopulation of neutrophils, first described in humans with systemic lupus erythematosus, [2] that may be separated from other granulocytes by discontinuous density gradient [2]. LDNs have a similar density to peripheral blood mononuclear cells (PBMCs) and are isolated alongside these cells, in contrast to normal density neutrophils (NDNs) which segregate with other polymorphonuclear (PMN) cells during density exclusion. [3,4]

LDNs have been described in pregnancy,[5] autoimmune disease,[6,7] cancer,[8,9] infection[10–13] and inflammation,[14,15] and speculated to contribute to pathophysiology. Across these studies, LDNs do not represent one discrete neutrophil sub-population, but rather a spectrum of multiple neutrophil phenotypes which vary in morphology, maturation and activation dependant on the underlying disease process. In addition, previously characterised monocyte-derived suppressor cells (MDSCs) that suppress T cell function can also contribute to the LDN population[16,17], further complicating the description of these cells between patient groups. Simply isolating LDN populations by density exclusion produces mixed populations of cells, and the lack of consistent LDN markers means that cell sorting techniques from whole blood are unsuitable.

Early studies counted LDNs based on density alone,[2] and in some cases LDNs are identified by flow cytometry as neutrophils in the PBMC layer. These studies are limited to characterising neutrophil phenotype without assessing the function of pure LDNs.[3,5,13,14,17–21] In addition, isolation of neutrophils within the PBMC and in the presence of monocytes can result in their activation. [22] For functional assays, further isolation of LDNs from PBMC is performed by FACS [10,12] which can also cause artefactual activation or by magnetic selection which is also problematic if the NDNs do not undergo the same protocol [7,23–27] and are instead subject to second density gradient or RBC lysis treatment. This is an important consideration as differences in isolation method can significantly alter granulocytes markers often measured in the context of LDN characterisation.[28] Furthermore many functional studies of LDNs have compared LDNs isolated from patient groups to healthy NDNs (not healthy LDNs) as a control.[15,29,30]

The described function of LDNs varies by the disease. In Systemic Lupus Erythematosus (SLE) and other autoimmune diseases, LDNs are highly inflammatory, produce type I interferon and activate T cells to produce TNFα and type II interferon.[7,25] A similar phenotype is also seen in some infections.[10] Conversely in *Mycobacterium tuberculosis* infection or following surgical stress, LDNs suppress T cell proliferation and IFNγ production.[26,27] The ability of LDNs to generate neutrophil extracellular traps (NETs) also varies between conditions. In SLE and Psoriasis, LDNs produce increased NETs compared with NDNs from the same donor,[14,23] while in Rheumatoid Arthritis, NET formation did not appear to differ,[30] but importantly healthy LDNs were not included as a comparator. Together, these data suggest that low neutrophil density may represent ‘disease specific’ neutrophil populations rather than a distinct class of neutrophils common to all conditions.

To interrogate whether LDNs are a distinct class of neutrophil with unique functional characteristics we first developed a robust method to isolate pure LDNs from healthy control blood. We then compared these to a disease control associated with infection and inflammation, namely Cystic Fibrosis, (CF). We have previously demonstrated a pro-survival phenotype and increased incidence of NET formation in CF neutrophils[31] but the contribution of LDNs to these findings has not been assessed.

In summary, we demonstrate a novel protocol to isolate highly pure populations of NDNs and LDNs from healthy donors. We define and comprehensively phenotype the characteristics of LDNs in the circulating neutrophil population of healthy donors and clinically stable CF patients, and found many similarities. We show that LDN nuclear morphology is less mature than autologous NDNs, and have reduced capacity for NET formation, despite similar levels of ROS generation and cell death by apoptosis. Finally, we demonstrate that NDNs can become low-density following activation with a number of inflammatory mediators. This research identifies an LDN population within healthy donors that have no significantly different function to circulating NDNs, aside from a perturbation of NET formation. In addition, it highlights the limitations of defining neutrophils by density alone and implies that neutrophils with low density are generated during inflammation or as a developmental step in maturation and are not a unique subset of cells with defined function.

## Results

### High purity LDNs and NDNs can be enriched from healthy donor blood by negative selection followed by Percoll density gradient

First, we compared LDN separation using the traditional method of a Percoll density gradient to a new strategy employing negative magnetic selection with the EasySep™ Direct Human Neutrophil Isolation Kit (STEMCELL), followed by discontinuous Percoll density gradient (Figures 1A and 1C). LDNs were distributed within a mixed population of lymphocytes and monocytes by the traditional isolation method (Figure 1B). Our new isolation strategy resulted in a highly pure (>95% by cytospin count) population of LDNs from healthy donors with no requirement for any further purification steps such as cell sorting (Figures 1D and 1E). LDNs isolated by this method are smaller and have segmented nuclear morphology in comparison to those found in the normal density layer, but lobes appear less defined and therefore more immature (Figure 1D). The median number of LDNs isolated by the new protocol from (N=10) healthy donors was 630,000 from 12 mls whole blood, which accounted for 3.62% of total neutrophils isolated on average (Figure 1F). This number is sufficient to perform functional assays.

**Figure 1:**
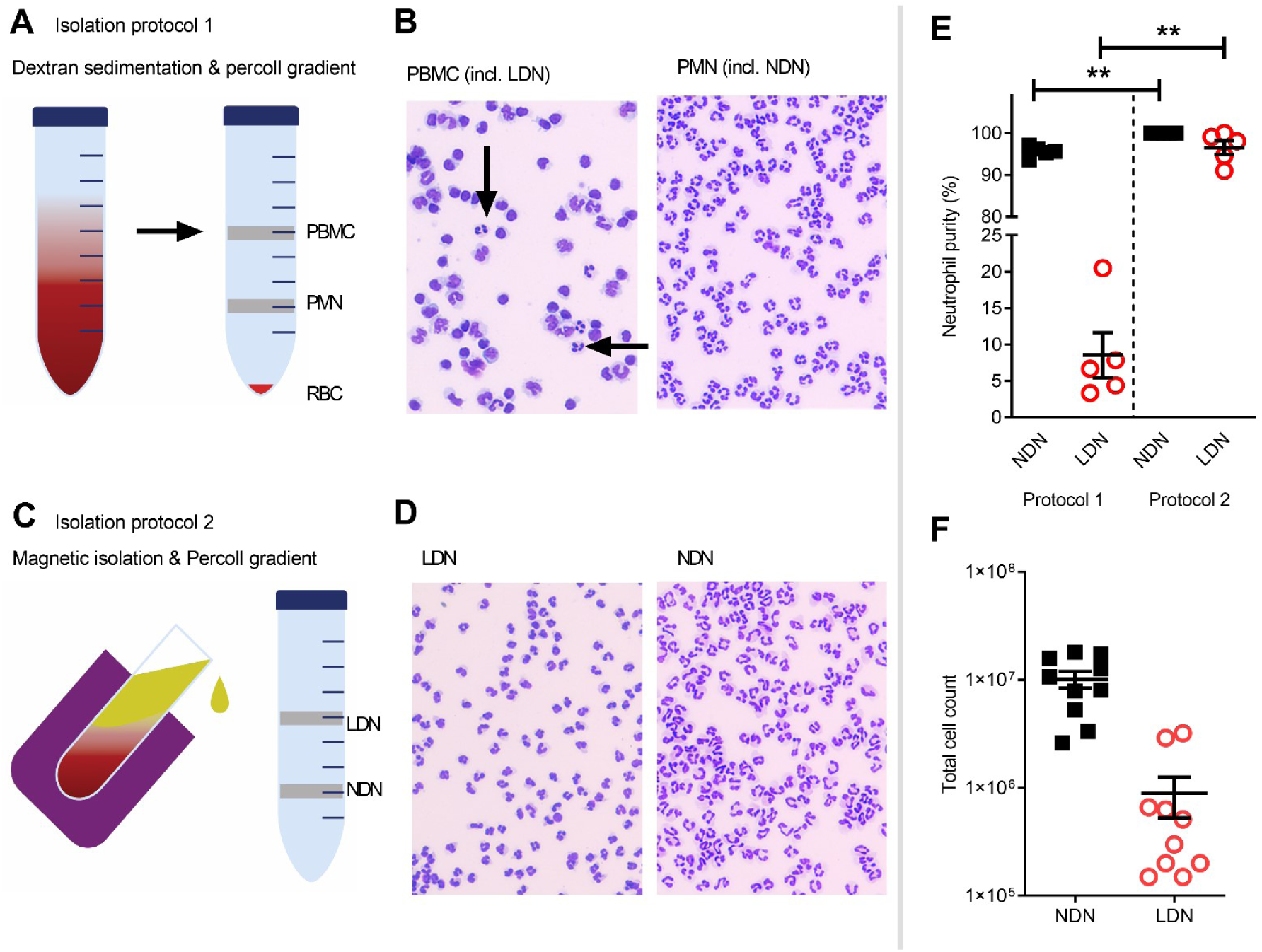
Isolation of a high purity population of low density neutrophils from healthy donors. **A)** Overview of traditional isolation protocol PBMC = peripheral blood mononuclear cells PMN = polymorphonuclear cells, RBC = red blood cells **B)** Representative images of cells isolated by traditional protocol, Black arrows show neutrophils in PBMC layer **C)** Overview of magnetic, negative isolation protocol, LDN = low density neutrophils, NDN= normal density neutrophils **D)** Representative images of enriched LDNs and NDNs isolated by magnetic, negative selection protocol **E)** Quantification of neutrophils isolated by each protocol, (N=5) **F)** Total number of neutrophils isolated from 12mls blood (N=10). Statistical testing with two tailed unpaired T test where **=p≤0.01 Data shows means ± SEM error bars.

### Healthy LDNs are indistinguishable from NDNs by flow cytometry

The markers used to identify LDNs differ between studies and between disease groups (literature reviewed in Table 1). These reports are frequently conflicting and a definitive set of LDN markers have yet to be resolved – in addition, very few studies have used healthy donor LDNs as the control group.

**Table 1:**
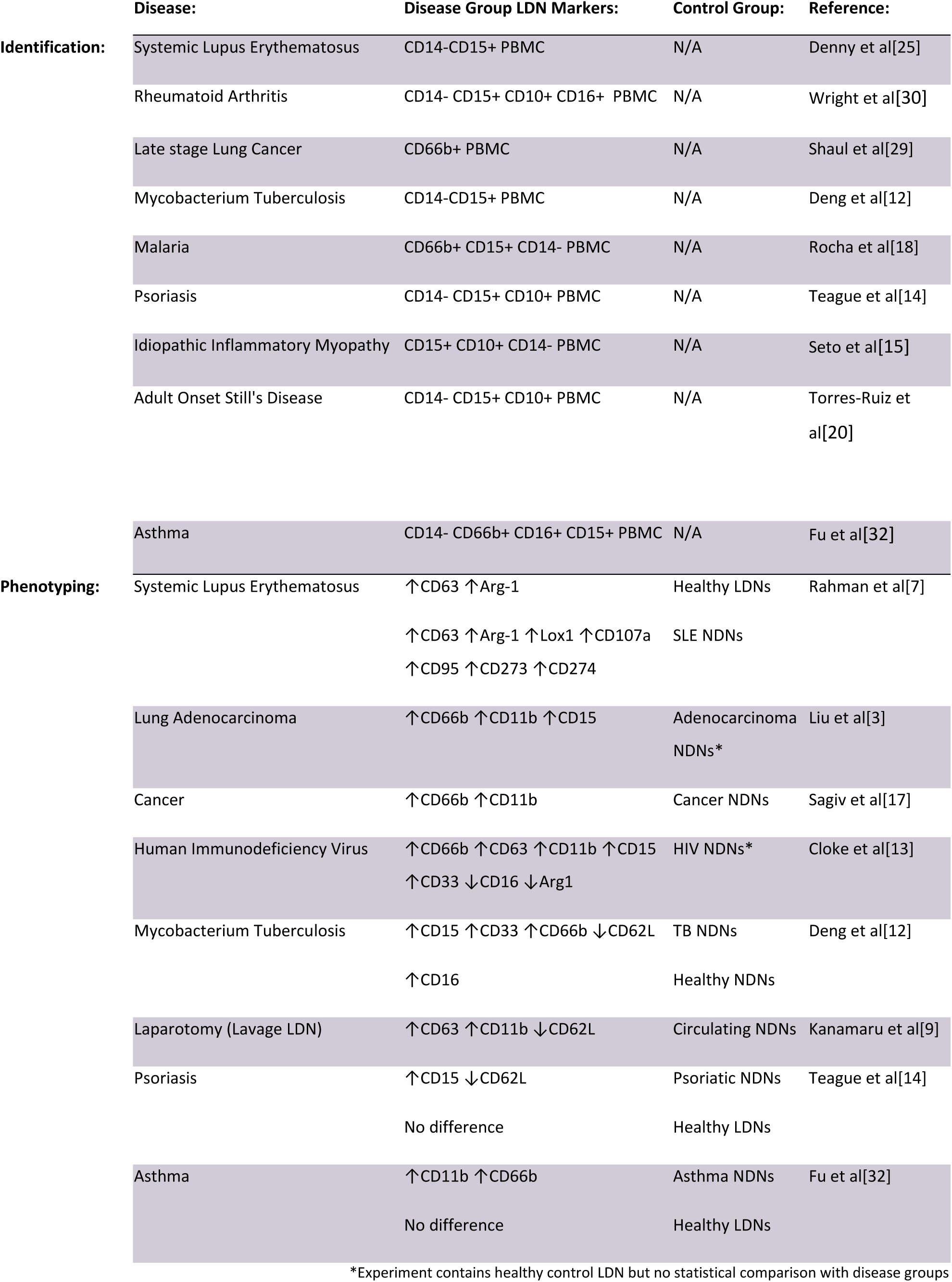
Overview of LDNs characterised by flow cytometry:

We have generated high purity LDNs and NDNs from healthy individuals in a method that minimises artefactual activation. We therefore used flow cytometry to characterise the LDN and NDNs isolated from healthy donors in an attempt to identify key markers of LDNs that may be used to identify these neutrophils without density separation, in a clinical setting for example.

First, we measured granulocyte maturation marker expression based on previously published observations. There was equal expression of CD66b, CD16, CD15 and CD10 on LDNs and NDNs isolated from healthy individuals by the new protocol, despite variation between donors (Figure 2B).

**Figure 2:**
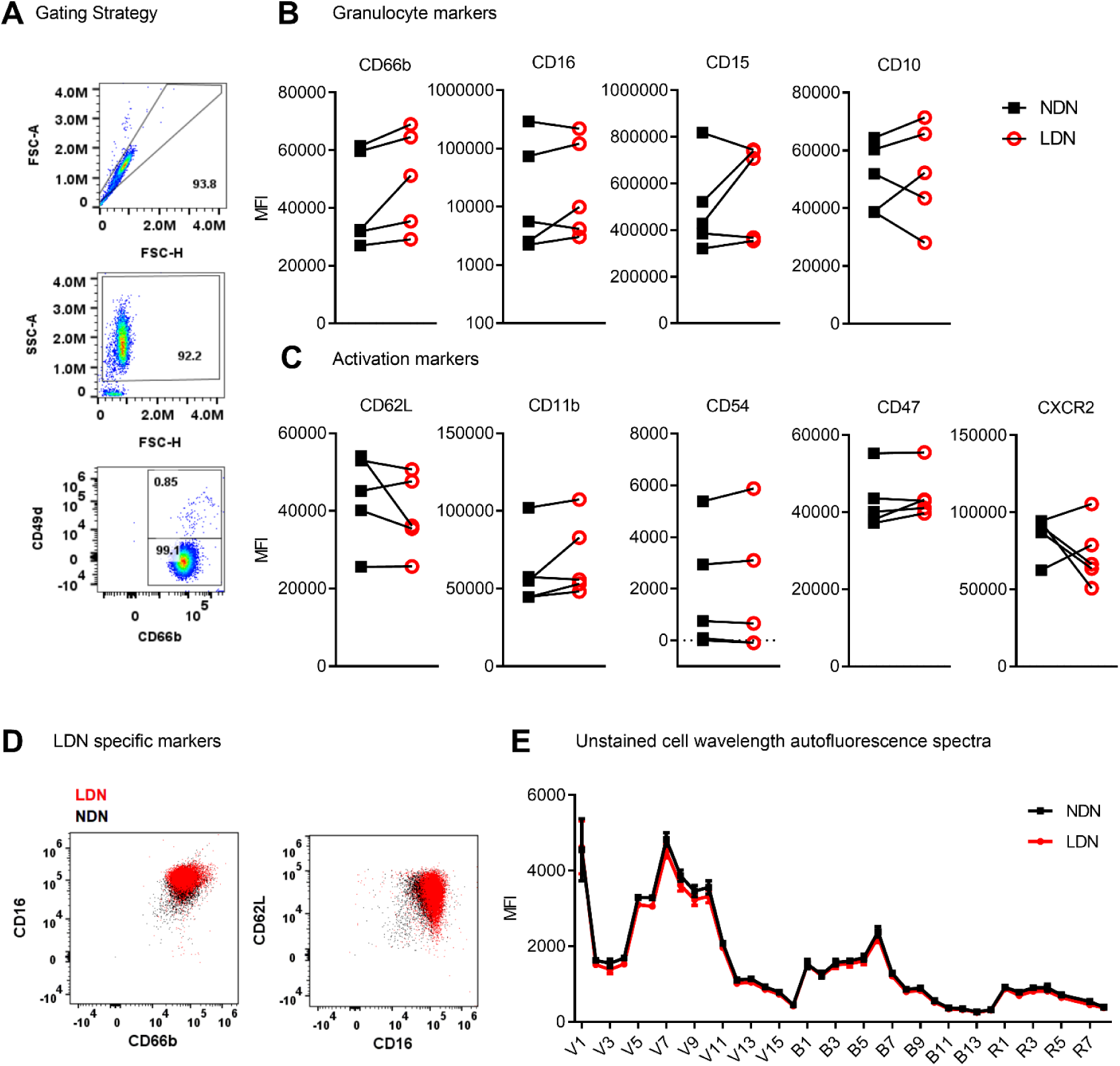
Expression of neutrophil markers on LDN and NDNs. **A)** Gating strategy to identify neutrophils (CD66b^+^ CD49d^-^) and remove any remaining eosinophils (CD66b^+^ CD49d^+^) after magnetic isolation. **B)** Markers of neutrophil maturation measured by flow cytometry, shown as mean fluorescence intensity (N=5). Linked samples show LDN and NDN from the same donor. **C)** Markers of neutrophil activation measured by flow cytometry, shown as mean fluorescence intensity (N=5). **D)** NDN overlaid with LDN from a representative healthy donor to demonstrate combinations of markers previously reported to identify low density neutrophils, CD16^HI^, CD66b^HI^ and CD62L^LO^,CD66b^+^ shown as mean fluorescence intensity. **D)** Spectral intensity recorded on unstained LDN and NDN cells (N=5, Mean ± SEM error bars).

Following this we measured markers of neutrophil activation, which have also been used to speculate on LDN function. Unlike previous studies in disease populations, L-selectin (CD62L) and integrin alpha M (CD11b) did not differentiate LDNs from NDNs (Figure 2C) in the same healthy individuals. We hypothesized that neutrophil migration markers may be different on LDNs. CD54 (I-CAM1) and CXCR2 expression also failed to differentiate between the two neutrophil populations, as did integrin associated protein (CD47) (Figure 2C). Furthermore, we were unable to show that LDNs were significantly phenotypically different to NDNs from the same donor, for any previously reported combinations of markers (Figure 2D). Finally, as different immune cells have characteristic auto-fluorescence and absorption of different wavelengths of light, a factor exploited by flow cytometry, we used a spectral flow cytometer to generate an auto-fluorescence spectrum for the isolated LDNs and NDNs and found no change in auto-fluorescence across all detectors (Figure 2E).

### LDNs show decreased NET formation but no difference in apoptosis or oxidative burst compared to NDNs

As marker expression is unaltered, we next asked if neutrophil function was different between LDNs and NDNs isolated from healthy donors. In both Systemic Lupus Erythematosus (SLE) and Idiopathic inflammatory myopathy, neutrophil extracellular trap (NET) formation is increased in LDNs when compared to disease group NDNs and healthy control neutrophils.[15,19] In these and other studies ‘control neutrophils’ comprise neutrophils found in the PMN layer, isolated by Ficoll/Percoll gradient NET formation by healthy donor LDNs is therefore poorly described (Reviewed in Table 2). Healthy LDNs produce fewer NETs following PMA stimulation than NDNs and therefore produce reduced fluorescence over the 6 hour time course (Figure 3A) although this did not reach statistical significance. We confirmed the presence of NETs in both LDN and NDN cultures with fluorescence microscopy and identified both ‘cloud-like’ DNA structures characteristic of NETs generated by PMA stimulation and some ‘stretched’ NETs more commonly seen *in vivo* (Figure 3B).

**Table 2:** Overview of NET formation in LDN populations:

| Disease: | NETs: | Reference: |
| --- | --- | --- |
| Systemic Lupus Erythematosus | ↑ Spontaneous NETs vs SLE NDNs and healthy NDNs<br>No difference upon PMA stimulation | Villanueva et al[23] |
| Rheumatoid Arthritis | No difference vs RA NDNs or healthy NDNs | Wright et al [30] |
| Mycobacterium Tuberculosis | ↑ Spontaneous NETs vs TB NDNs<br>↓ PMA stimulated NETs vs TB NDNs | Su et al[24] |
| Laparotomy (Lavage) | Post-operative LDNs produce NETs spontaneously | Kanamaru et al[9] |
| Abdominal surgery (Blood) | Circulating LDNs produce NETs spontaneously | Kumagai et al[27] |
| Psoriasis | ↑ NETs vs Psoriatic NDNs | Teague et al[14] |
| Idiopathic Inflammatory Myopathy | ↑ Spontaneous NETs vs IIM NDNs<br>(Both are higher than healthy NDNs) | Seto et al[15] |
| Adult Onset Still's Disease | ↑DNA-elastase complexes correlates with LDNs<br>AOSD serum ↑ NETs | Torres-Ruiz et al[20] |
| Equine Asthma | ↑ NETs vs Equine Asthma NDNs<br>↓ NETs vs Healthy Equine LDNs | Herteman et al[33] |

**Figure 3:**
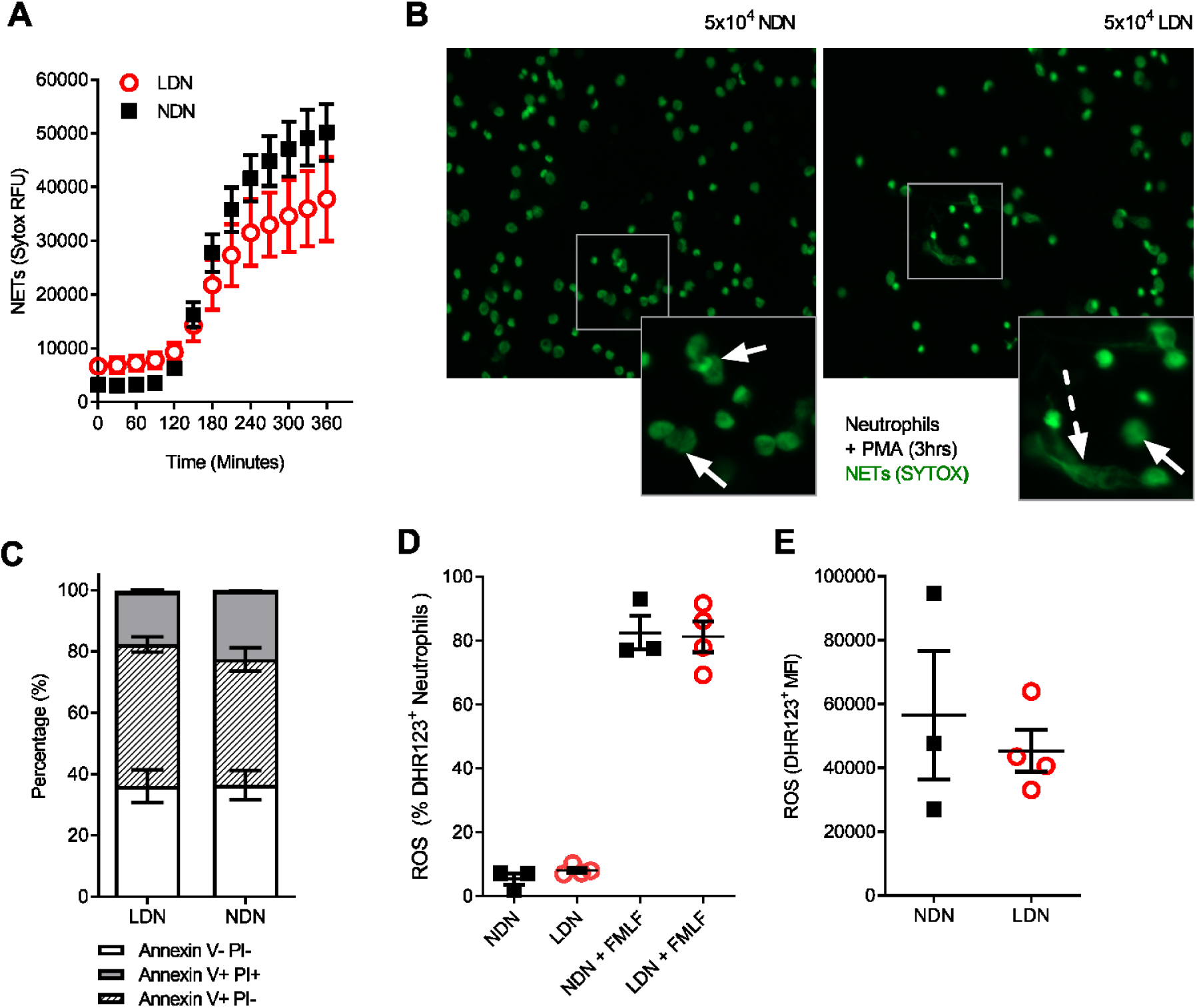
LDNs form fewer NETs than NDNs in healthy individuals. **A)** NET formation by 5×10^4^ LDNs or NDNs, stimulated with PMA measured by SYTOX green fluorescence over 6 hours (N=4-6). **B)** Representative images of NETs formed by LDNs and NDNs after 3 hour stimulation with PMA. Solid arrows show ‘cloud-like’ NETs and dashed arrows show ‘stretched’ NETs. **C)** Apoptotic neutrophils measured by Annexin V and PI expression after 20 hours in culture (N=4-5) **D)** Percentage of LDNs and NDNs positive for ROS before and after FMLF stimulation, measured by DHR123+ fluorescence. (N=3-4) **E)** ROS generated by LDNs and NDNs in response to FMLF stimulation, expressed as DHR123 mean fluorescence intensity (N=3-4).

Neutrophils would normally undergo apoptotic cell death before clearance from the site of inflammation. We have previously shown in patients with CF that extended neutrophil life span contributes to increased NET formation.[31] In Rheumatoid Arthritis (RA), LDNs have reduced apoptosis compared with RA NDNs, even when treated with GM-CSF.[30] We therefore cultured isolated LDNs and NDNs for 20 hours *in vitro* then stained with Annexin V and Propidium Iodide to determine the levels of apoptosis. There was no significant difference in apoptosis rates between the neutrophil densities, with ∼70% of LDNs and NDNs undergoing apoptosis at the 20 hour time point (Figure 3C). In SLE, LDNs produce similar amounts of ROS to NDNs[25] yet production of reactive oxygen species (ROS) is required for some types of NET formation[34] and we have shown differences in NET formation in healthy LDNs. We therefore questioned if ROS production is impaired in healthy LDNs. LDNs isolated from healthy donors are capable of producing ROS (Figure 3D) and the extent of ROS production is the same as that in NDNs upon FMLF stimulation (Figure 3E).

### T cells co-cultured with LDNs and NDNs have similar proliferation responses and IFN-γ production

Neutrophil - T cell interaction is now well described as a mechanism of innate and adaptive immune cross talk. Sub-populations of neutrophils have been shown to suppress proliferation of T cells[35] and LDN responses are often linked to activation or suppression of T cell responses, particularly in the tumour microenvironment.[36,37] Neutrophils in circulation suppress T cell proliferation and T cells stimulated with BCG, have decreased proliferative responses and IFN-γ production in the presence of LDNs.[26] Progenitor cells from the bone marrow however are incapable of T cell suppression[38] and while LDNs from SLE patients activate T cell IFN-γ and TNF production they do not affect proliferation.[7] To determine how the LDNs found in healthy controls compare with these phenotypes, we co-cultured LDNs and NDNs from healthy donors for 96 hours with T cells. Beads coated with anti-CD3 / anti-CD28 antibodies were used to activate the T cells. Total CD4^+^ or CD8^+^ T cell numbers were unaltered by co-culture with neutrophils for 96 hours (Figures 4B and 4F). The proportion of CD4^+^ and CD8^+^ T cells producing IFN-γ after 96 hour co-culture with LDNs or NDNs was also unaltered compared with T cell only controls (Figures 4C and 4G) and amount of IFN-γ was unchanged (Figures 4D and 4H). Proliferation was measured by dilution of CFSE labelled T cells and quantified by flow cytometry. T cells cultured alone showed significant proliferation following 96 hour stimulation (Figures 4E and I). Co-culture with either LDN or NDNs resulted in decreased numbers of proliferating CD4^+^ and CD8^+^ T cells, but this response was not significantly different between the two neutrophil densities. (Figures 4E and 4I respectively).

**Figure 4:**
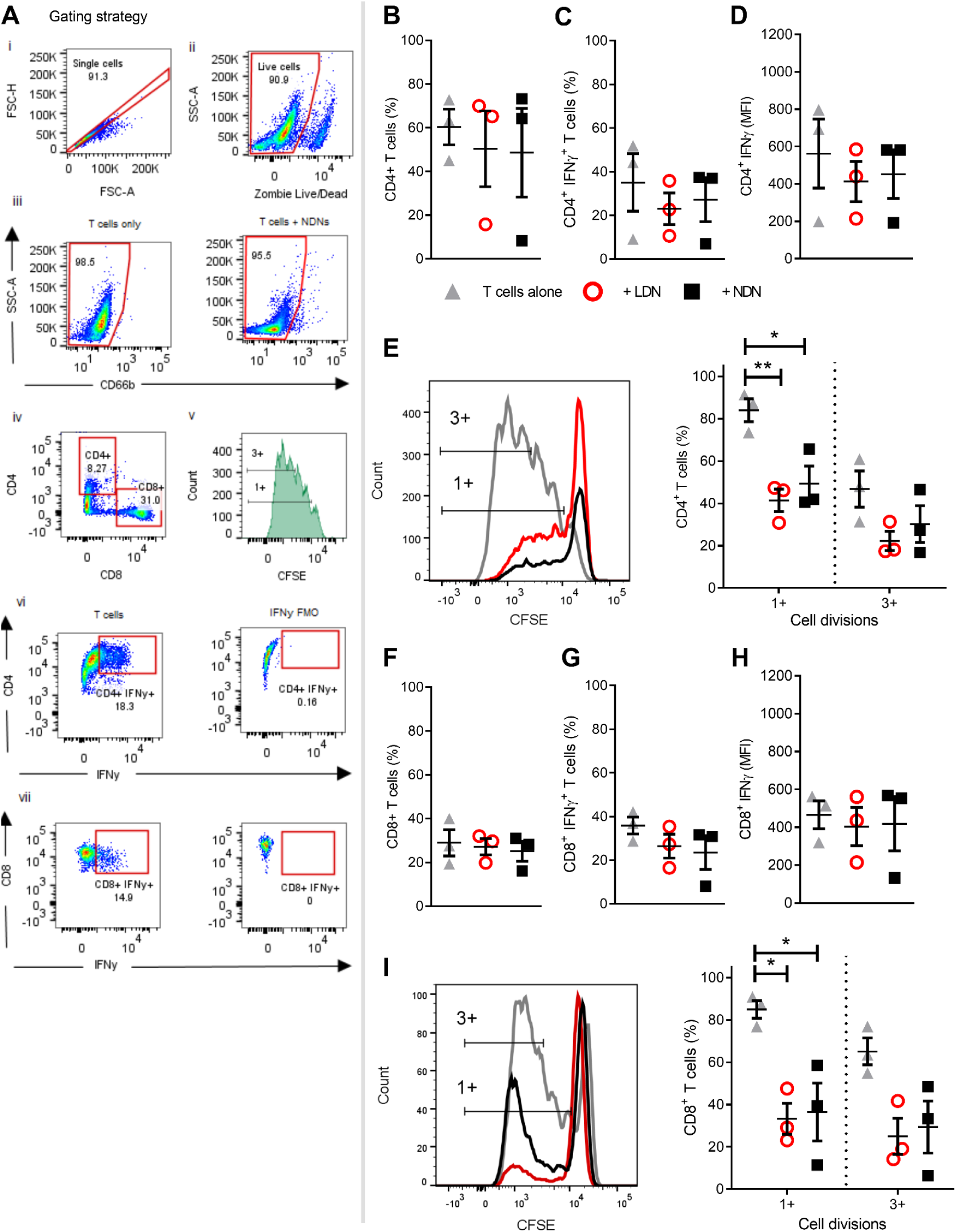
Co-culture of T cells with either LDNs or NDNs reduces T cell proliferation. **A)** Gating strategy to identify T cells after 96 hour co-culture with neutrophils. **B)** Percentage of CD4^+^ T cells in culture after 96 hours (N=3). **C)** Percentage of CD4^+^ T cells producing intracellular IFN-γ after 96 hour co-culture with neutrophils then 4 hour stimulation with T cell activation cocktail (N=3). **D)** Amount of intracellular IFN-γ production in CD4^+^ T cells, shown as mean fluorescence intensity (N=3). **E)** CD4^+^ T cell proliferation, measured by reduction in peaks of CFSE fluorescence (N=3). **F)** Percentage of CD8^+^ T cells in culture after 96 hours (N=3). **G)** Percentage of CD8^+^ T cells producing intracellular IFN-γ after 96 hour co-culture with neutrophils then 4 hour stimulation with T cell activation cocktail (N=3). **H)** Amount of intracellular IFN-γ production in CD8^+^ T cells, shown as mean fluorescence intensity (N-3). **I)** CD8^+^ T cell proliferation, measured by reduction in peaks of CFSE fluorescence (N=3). All data represented as individual values with mean ± SEM). Statistical testing by one way ANOVA with multiple comparisons.

### Healthy NDNs can be induced to form LDNs upon activation with TNF, LPS or FMLF

The generation of LDNs is poorly understood, yet increased numbers are observed in many inflammatory environments.[12,15] We therefore hypothesised that activation during various disease states would affect the production of LDNs from already circulating NDNs. Neutrophils were isolated from healthy donor whole blood by dextran sedimentation followed by traditional discontinuous Percoll density gradient (55% / 70% / 81%). All neutrophils recovered in the ‘PMN layer’ (NDNs) were then cultured either untreated or in the presence of three different inflammatory stimuli (TNF or LPS for 2 hours, or FMLF for 30 minutes), or left untreated. Following stimulation, the neutrophils were again separated by discontinuous density gradient as before (Figure 5A). NDNs which were untreated with stimuli and re-run through the Percoll gradients were largely recovered in the NDN layer as before (Figure 5B), indicating that the isolation was reproducible and that two hours’ incubation at 37°c had not altered their density. However, NDNs activated with any of the inflammatory stimuli were now predominantly recovered from the LDN phase (Figures 5B and 5C). Accompanying cytospins of LDNs from each treatment conditions demonstrate the varied neutrophil morphologies (Figure 5D). The nuclear structure in the primed and activated cells retains its mature, segmented phenotype, but the cytosol in fMLF treated cells became vacuolated and LPS treated neutrophils demonstrated membrane ruffling and protrusions from the cell surface. Control cells that were isolated as LDNs after 2 hour culture had a similar appearance to NDNs before culture.

**Figure 5:**
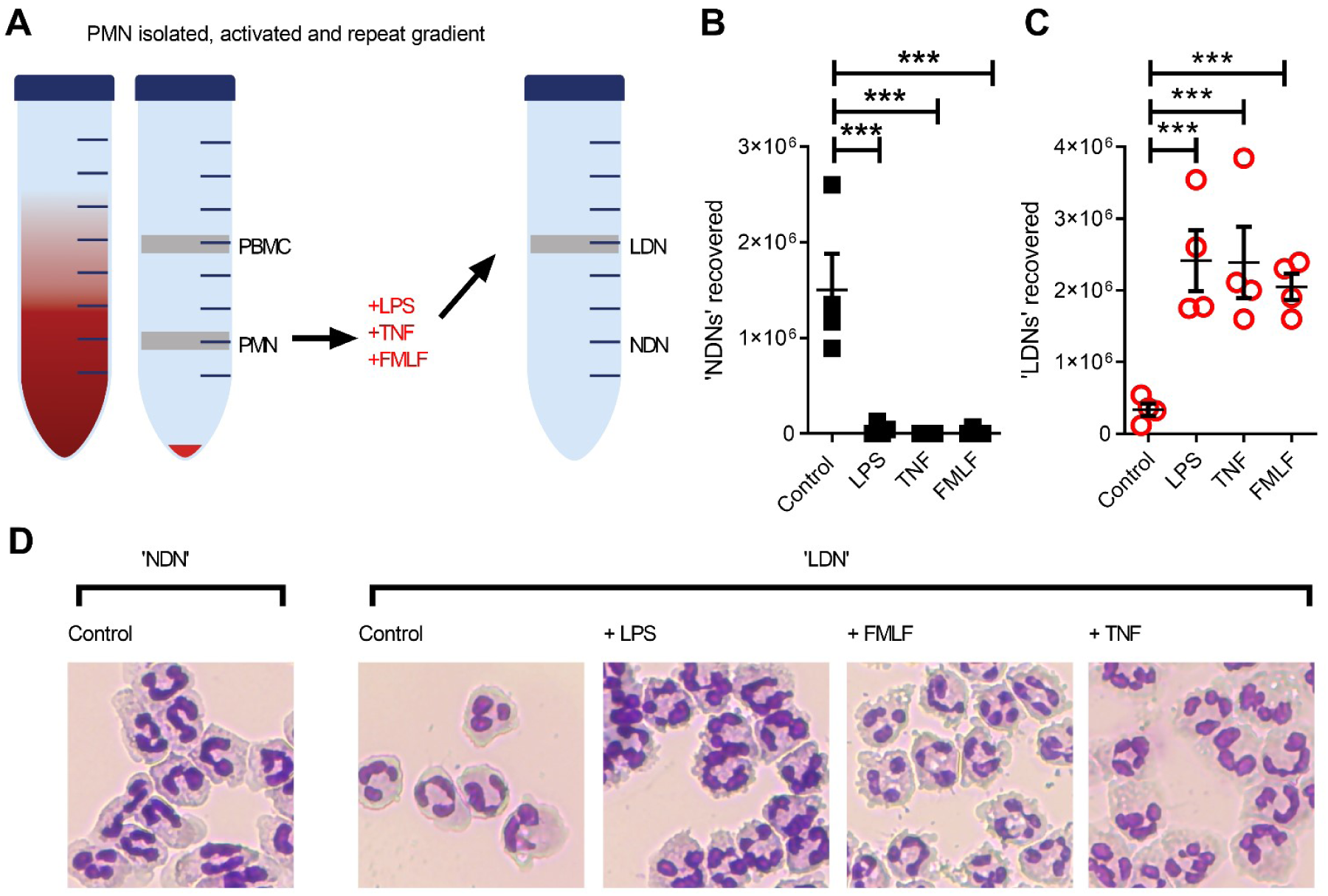
NDNs become LDNs upon activation with inflammatory stimuli. **A)** Diagram of method. PMN isolated by traditional methods are activated *in vitro* with inflammatory stimuli and then undergo repeat Percoll gradient. **B)** Number of neutrophils recovered from the ‘PMN/NDN’ layer after second Percoll gradient. (N=4) **C)** Number of neutrophils recovered from the ‘PBMC/LDN’ layer after second Percoll gradient. (N=4) **D)** Representative cytospin images of neutrophils recovered in each condition. Statistical analysis by One-way ANOVA with multiple comparisons where ***=p≤0.001.

### NDNs but not LDNs are significantly elevated in the circulation of people with Cystic Fibrosis

We have shown that inflammation or activation of healthy NDNs can generate LDNs. We therefore asked if we could isolate LDNs with our new protocol from a patient cohort with a well characterised neutrophil phenotype. Our lab has previously demonstrated a pro-survival, apoptosis resistant response in neutrophils from people with Cystic Fibrosis that results in increased production of NETs.[31] We asked if LDNs accounted for the significant increase in neutrophils seen in this patient group. The median number of LDNs isolated from people with CF was 230,000 (N=5) and accounted for 4.64% of total neutrophils. This is not significantly different to the proportion found in healthy individuals (described in Figure 1). The total number of NDNs however, was significantly higher in people with CF when compared to healthy controls (p=0.0047, by two tailed T test, Figures 1D and 6A). In addition the pro-survival ‘anti-apoptotic’ phenotype previously described in CF neutrophils was present in both CF NDNs and LDNs (Figure 6B). The mean number of live cells after 20 hour culture was 45.23% ± 10.43% (Mean ± SEM) for CF NDNs and 53.18% ± 10.26% for CF LDNs compared with 36.03% ± 5.31% for healthy LDNs and 36.42% ± 4.81% for healthy NDNs. NET formation is reduced in CF LDNs compared with CF NDNs, mirroring the phenotype observed in healthy individuals (Figure 6C).

**Figure 6:**
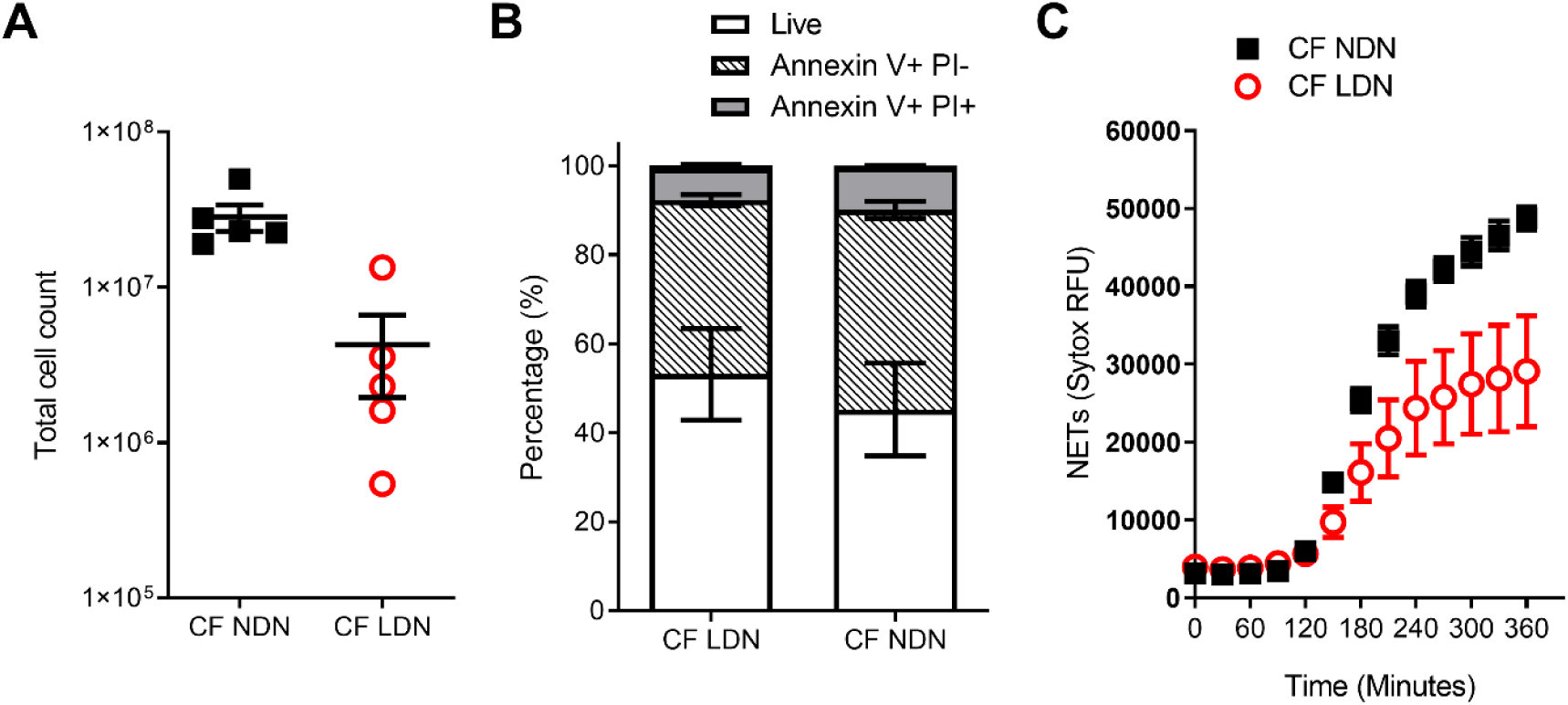
LDNs are not elevated in people with Cystic Fibrosis. **A)** Total numbers of LDNs isolated from 12 mls blood by magnetic separation and Percoll gradient. (N=5) **B)** Rate of apoptosis after in vitro culture for 20 hrs, measured by Annexin V and Propidium Iodide staining. (N=4-6) **C)** NET formation, measured by SYTOX green fluorescence over 6 hours after stimulation with 100nM PMA (N=5-6) Data shown as mean ± SEM error bars.

## Discussion

We have developed a new protocol to isolate LDNs from whole blood that can be used to study LDN phenotype and function in healthy controls and disease populations. We show that healthy LDNs are present in circulation, have an immature segmented nuclear morphology and, in contrast to LDNs observed in SLE, have reduced capacity to form NETs - yet lack any distinguishing markers of activation or maturation characteristic of LDNs in disease. Healthy LDNs have unaltered ROS production, undergo apoptosis at the same rate as autologous NDNs, and furthermore do not alter T cell proliferation or IFNγ production in comparison to NDNs.

Our simple protocol, of magnetic negative selection of total neutrophils from small volumes (12 mls) of blood followed by Percoll gradient, isolates highly enriched populations of low and normal density neutrophils from healthy donors. This is rapid, less likely to activate neutrophils than methods such as FACS, effectively nullifies any chance of activation by monocytes, and leads to >95% pure populations of LDN. Healthy control LDNs are usually not quantified in studies of LDN function during disease or infection owing to the perception that they cannot be readily harvested. Previous studies therefore lack the crucial control of healthy LDNs.

Phenotypic markers used to identify LDNs differ between studies and disease groups and as such a definitive set of LDN markers has yet to be resolved (Table 1). Granulocyte markers, CD66b, CD15 and CD16 (FcγRIII) are frequently used to identify LDNs within the PBMC cell population (see Table 1). In Rheumatoid Arthritis and ANCA positive Vasculitis, CD16 expression is reduced on LDNs[30,39] while in lung adenocarcinoma and HIV infection, CD15 is elevated on LDNs.[3,13] Furthermore LDNs in gastric cancers have high CD66b expression.[9] CD10 (Neprilysin, CALLA) expression is increased on mature neutrophils. These are discernible from CD10^LOW^ immature neutrophils which suppress T cell activation.[40] In severely infected patients, CD10^LOW^ neutrophils increase with infection severity and this is correlated with increased numbers of LDNs.[11] Surprisingly we found that none of these markers effectively differentiated healthy LDNs from healthy NDNs, suggesting that the differences in many published studies are not intrinsic features of less dense neutrophils.

We therefore turned our attention to the function of LDNs. Healthy LDNs had no differences in rate of apoptosis or oxidative burst compared to NDNs. LDNs from healthy individuals produced fewer NETs than NDN, in direct contrast to LDNs isolated from chronic inflammatory conditions such as psoriasis and idiopathic inflammatory myopathy, where LDNs produce increased numbers of NETs.[14,15] It is difficult to reconcile many of the features of LDNs as healthy LDNs have normal expression of CD10 and CD16 characteristic of mature neutrophils and no reduction in ROS production in spite of having an immature nuclear morphology.

A major function attributed to LDNs is their ability to either activate T cells as in SLE,[7] or to suppress T cell proliferation as described in cancer.[27] Our data show that in healthy donors, LDNs and NDNs did not affect intracellular IFNγ or frequency of CD4^+^ and CD8^+^ T cells, but both populations significantly impaired T cell proliferation. T cell suppression assays are not without limitations, as neutrophils may prevent αCD3/αCD28 bead stimulation of T cells in some part by phagocytosing beads.[41] Furthermore, after 96 hours in culture, ‘suppression’ of T cell division may be due to the presence of apoptotic cells. We demonstrated no significant differences between LDNs and NDNs on T cell responses *in vitro* in spite of these limitations. Absence of CD10 expression is a marker used in characterisation of immature LDNs that can suppress T cells[11] yet the relationship between CD10 and LDNs is complex. LDNs in autoimmune disease are CD10^+^[25] and LDNs identified in cardiovascular risk associated with psoriasis are identified by CD10^hi^ expression.[14] As CD10 expression was equal in LDNs and NDNs from healthy individuals and T cell suppression was seen in both neutrophil densities, we would propose that the T cell suppression is disease specific, and perhaps CD10^−^ LDNs seen in disease may be a result of emergency granulopoesis[40] and are therefore not present in healthy individuals.

*Mycobacterium tuberculosis (Mtb)* infection of healthy blood samples induces additional numbers of LDNs,[12] suggesting that LDNs may be directly generated by inflammation or infection. To assess if we could induce LDNs from NDNs, we isolated NDN and then stimulated cells with either LPS, TNF or FMLF before separating once again by Percoll gradient (Figure 5). Activated, previously normal density neutrophils now segregated to the PBMC/LDN layer. While this does not explain the population of LDNs isolated from healthy donors at baseline, this experiment suggests that neutrophil priming and stimulation in disease could be an important driver of increased LDN populations.

CF is a progressive disease associated with neutrophilic lung pathology. We have previously demonstrated that neutrophils in CF have a pro-survival phenotype[31] but it is unknown whether LDNs contribute to this observation. Interestingly, people with CF had significantly increased levels of NDNs in comparison to healthy donors, but a similar proportion of LDNs, suggesting they are not a major feature of the neutrophilia seen in CF. Notably, LDNs in our CF population were also resistant to apoptosis mirroring the phenotype seen in CF NDNs, yet were less capable of NET formation than CF NDNs (Figure 6C). These data may be explained by the compartmentalised nature of inflammation in the CF lung while systemic inflammation is less prominent in times of disease stability.

Low density neutrophils have been given significance in diseases ranging from cancer to autoimmune disease. Our data demonstrate that LDNs are part of the normal spectrum of neutrophils and that in health, aside from a reduction in NET formation capacity, they do not display a divergent phenotype. Furthermore, known activators of neutrophils can induce NDNs to become LDNs. We propose that standardised density protocols for neutrophil and mononuclear cell isolation has incorrectly led to the characterisation of LDNs a highly unique sub-population of neutrophils with defined function. Low neutrophil density could actually be a reflection of maturation or activation status. It may also be suggested that in health and disease that neutrophils range in sizes and densities, and the application of arbitrary cut-offs particularly in states of inflammation may have led to an overestimation of the intrinsic functions of LDNs.

## Methods

### Ethical approval, blood collection and serum separation

Healthy blood was collected in accordance with The Centre for Inflammation Research Blood Resource (AMREC, 148 15/HV/013). Clinically stable CF patients with one F508del mutation were recruited according to NRS Bioresource, East of Scotland research ethics committee 15/ES/0094.

12 ml of blood was collected into a tube containing 3.4% sodium citrate to prevent coagulation. Serum was isolated by centrifugation at 350 x g for 20 minutes at RT.

### NDN and LDN isolation and quantification

Total neutrophils were isolated from 12ml healthy or CF patient blood by negative selection using an EasySep™ Direct Human Neutrophil Isolation Kit (Stem Cell). LDN and NDNs were then isolated by discontinuous Percoll density gradient. Briefly, 27ml Percoll (GE Healthcare) was added to 3ml 10x PBS^-/-^ (Gibco) to create a 100% solution. Then, 3ml 81% Percoll diluted in 1x PBS^-/-^ (Gibco) was added to a 15ml tube (Falcon) carefully followed by a 3ml layer of 70% Percoll and finally, total neutrophils, resuspended in 3ml 55% Percoll. Centifugation at 720xg without brake was performed at RT for 30 minutes. NDNs were recovered from the 71%/55% interface and LDNs from 70%/81% interface (see Figure 1A). Cell counts were performed on a NucleoCounter® NC-100 (Chemometec). Cytospins were performed in 1% BSA (Sigma) in PBS^-/-^ in a Cytospin 4 (Thermo) and stained with Kwik-Diff stains (Shandon) according to manufacturer’s instructions.

### Flow cytometry and auto-fluorescence spectral map

1×10^6^ LDN or NDN were suspended in PBS^-/-^ + 2% fetal calf serum (Gibco) and stained with CD66b, CD16, CD15, CD10, CD11b, CD54, CD47, CXCR2 and CD62L antibodies (Biolegend) for 30 mins at 4°c. Samples were then analysed on an Aurora Spectral Flow cytometer (Cytek). Gating strategy is shown in figures. Auto-fluorescence was measured by mean fluorescence intensity in each detector on unstained cells to produce a spectral map.

### Apoptosis

5×10^5^ LDN or NDN were suspended in 200μl PBS^-/-^ +2% donor serum and incubated at 37°c, 5% CO_2_ for 24 hrs. Neutrophils were then stained with Annexin-V-FLUOS Staining Kit (Sigma) following manufacturer’s instructions and analysed by flow cytometry on an LSR Fortessa (BD) and FlowJo software (BD) based on previously described methods.[42]

### NET formation

5×10^4^ LDN or NDN were suspended in 200μl PBS^-/-^ +2% donor serum. NET formation was stimulated with 100nM PMA and detected with SYTOX™ Green nucleic acid stain (Thermo) at 528nm during 6 hr incubation at 37°c in a Synergy HT (BioTek) plate reader (as previously described).[31] NET formation was confirmed by fluorescence microscopy.

### Reactive oxygen species

2.5×10^5^ LDN or NDN were suspended in 500μl HBSS^+/+^ (Gibco) and stained with Dihydrorhodamine 123 (Thermo) as previously described[43] followed by stimulation with 10nM fMLF for 20 minutes at 37°c. The assay was stopped by transference to ice for 10 minutes before fixation with 4% PFA. Cells were analysed by flow cytometry. DHR 123 is measured by 488 laser excitation with a 525/50 filter.

### T cell & Neutrophil co-culture

1×10^5^ T cells were stained with CFSE (Thermo) and cultured in 200ul RPMI (Gibco) + 10% HI-FCS (Gibco) + 1% Peniciliin/Streptomycin (100x, Gibco) + 1% L –glutamine (100x, Gibco) with 2×10^4^ Human T-Activator CD3/CD28 Dynabeads (Thermo) alone or with 2×10^5^ LDNs or NDNs for 96 hours at 37°c 5% CO_2._ Activation cocktail containing protein transport inhibitors (500x eBioscience) was added for 4 hours then stained with Live/Dead Yellow (Thermo), followed by antibodies for CD4, CD8, CD66b (Biolegend) and intracellular IFNγ (Biolegend) following fixation and permeabilisation with 1x Fix/Perm solution (BD). Cells were then run on an LSR Fortessa flow cytometer (BD) and analysed with FlowJo Software (BD). Co-culture conditions were adapted from previously described methods.[35]

### Neutrophil activation

5×10^6^ PMN were suspended in 1ml HBSS^-/-^ (Gibco) alone or with either 100ng/ml LPS (Invivogen), 100 ng/ml TNF (R&D) for 2 hours at 37°c, 5% CO_2_, with gentle agitation at 1 hour. For FMLF treatment, 10nM fMLF (Sigma) was added after 90 minutes for the remaining 30 minutes of incubation. Neutrophils were then washed x2 in HBSS^-/-^ and resuspended in 55% Percoll for repeat density gradient.

## Statistics

Where two groups were analysed a two tailed, unpaired T test was used, where 3 or more groups were analysed, One-way analysis of variance was used with multiple comparisons. All graphs show mean ± standard error of the mean as average and error bars. Data was analysed in GraphPad Prism 8.

## Acknowledgements

We would like to acknowledge the contributions of Professor Julia R Dorin, University of Edinburgh to the discussion and development of this work. Flow cytometry data was generated with support from the QMRI Flow Cytometry and cell sorting facility, University of Edinburgh. This work was funded by a Chief Scientist Office Senior Clinical Academic Fellowship (SCAF/16/02) awarded to RDG and Medical Research Council Senior Fellowship (G1002046) awarded to DJD.

